# EzReverse – a Web Application for Background Adjustment of Color Images

**DOI:** 10.1101/2024.05.27.594095

**Authors:** Xinwei Song, Joachim Goedhart

## Abstract

Fluorescence imaging is extensively utilized in biological research to study biomolecules at the single-cell level. Fluorescence images have a black background due to the inherent nature of fluorescence imaging. The black background fits well with dark themes that are used on screens and presentations. On the other hand, inverting the color images to display the signals on a white background can aid the human visual system in capturing detailed information or subtle features. In addition, white backgrounds match better with printed media. However, current methods to invert or change the background of color images require multiple steps or do not offer flexibility in the adjustment which is important when dealing with complex images without raw data. To facilitate easy access to background inversion and flexible modifications, ezReverse was developed. Two methods are implemented, the first based on color space transformation followed by inverting the lightness, while the second method identifies grayscale values which are subsequently modified. The web app ezReverse is a versatile tool, that accommodates multiple color spaces, kernel filters, and gamma correction for optimal inversion of color images. It is hosted on the Shiny Python platform and can be conveniently accessed online: https://amsterdamstudygroup.shinyapps.io/ezreverse/

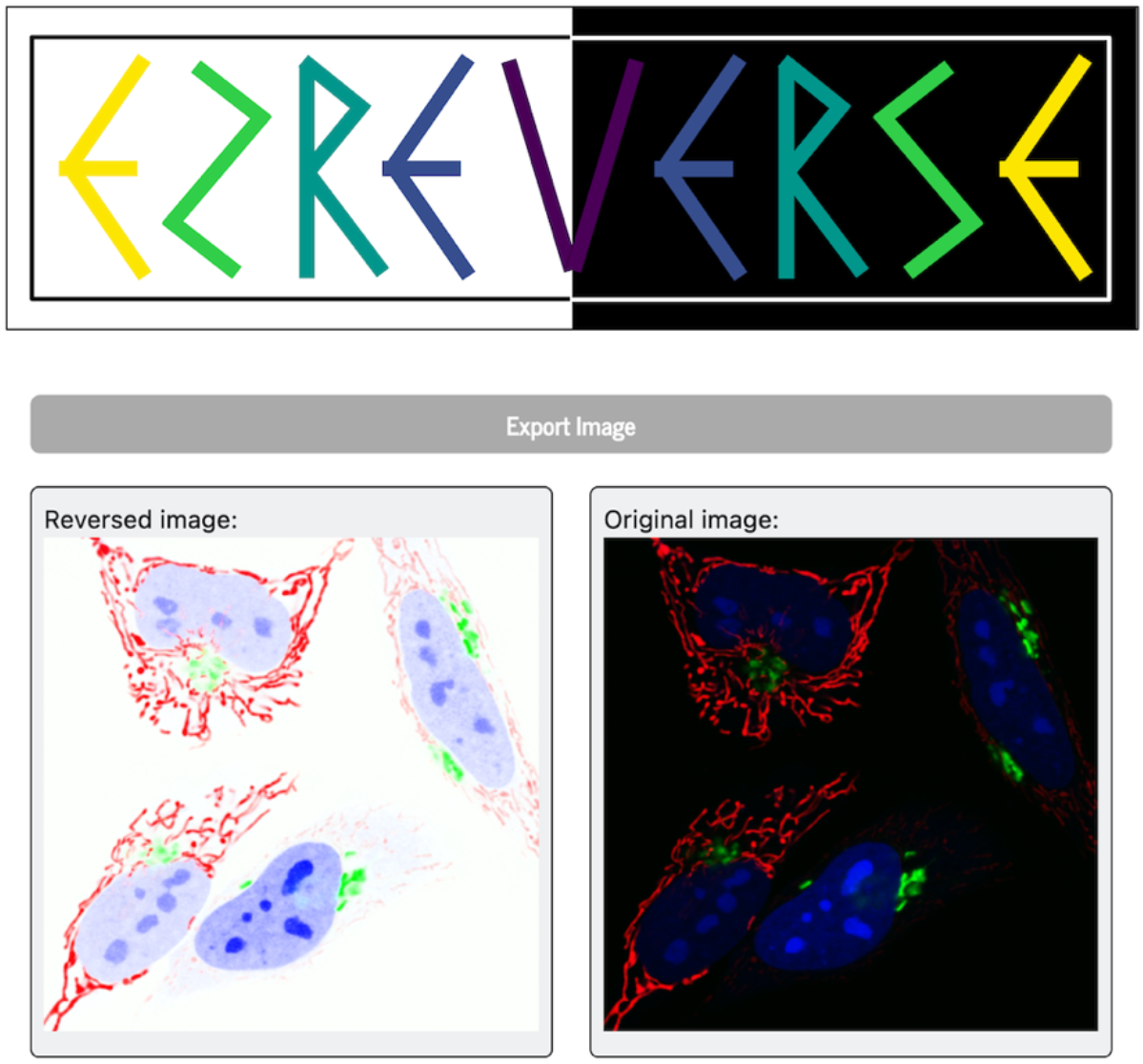

## Introduction

Fluorescence imaging has become a key technology for monitoring protein dynamics, biomolecules, biochemical alterations, and cellular states. This technology allows for the characterization of the stochasticity and heterogeneity that biological systems exhibit [1]. Nowadays, a wide variety of fluorescence microscopy techniques is available that all use fluorescent substances that provide contrast [2]. The intrinsic nature of fluorescence creates images where the fluorescent molecules produce a bright signal against a dark background. The high contrast is obtained since we detect light from the emitted fluorescence, but the background, where fluorescence molecules are absent, remains dark. This stark contrast can make it challenging for the human visual system to discern the finer details within the image.

Reversing such images has a broad range of benefits. Reversing black background images, particularly those with dim features, not only enhances perceptual contrast, thereby making the fine structure and image information more easily captured by the human visual system [3], but also augments the detection and assessment in some disease imaging. Following image inversion, even a single white spot, which may not be apparent on a standard black image, becomes readily recognizable as a black spot on a negative image. Previous studies have shown that such inversion improves mammographic evaluation, facilitating the detection of small yet crucial signs of breast cancer such as microcalcifications [4].Utilizing both the original and reversed images simultaneously has been found to enhance diagnostic sensitivity and accuracy among less biased readers, like junior EM residents or medical students, when compared to conventional images [5].

The black background of fluorescence images may also clash with light backgrounds that are common in printed media. Inverting fluorescence images for publications may be a better choice. On the other hand, standard fluorescence images fit well with slides that have a black background. Yet in this situation the plots and other elements that have a white background may be a poor match and inversion of these elements may result in more aesthetically pleasing slides.

Inverting grayscale images is a straightforward process since every pixel is encoded by a single value. In contrast, color information is encoded by multiple parameters, typically consisting of three channels (or images). The most ubiquitous format for such color representation is the RGB format that uses three coordinates to encode the red, green and blue component, which are associated with the sensitivity of cones in the human visual system. The RGB color model is the basis for many display technologies, as well as digital imaging and video, making it a fundamental color space in the digital world.

The combination of these three primary color channels produces a color image. When one inverts a color image in RGB colorspace by inverting each R, G, and B component, this inverts black, white and gray values as expected. However, inverting an RGB image changes the color. For instance, blue (0,0,255 in R,G,B code) turns into yellow (255,255,0, in R,G,B code) when inverted.

The HSL color model [6] is organized around a cylindrical-coordinate system, where each color is represented by three components: Hue (H), Saturation (S) and Lightness (L). Hue denotes the type of color and is measured in degrees (0 to 360). In this system, red is at 0°, green at 120°, and blue at 240°. Saturation is expressed as a percentage, indicating the intensity or purity of the color. Lightness, also expressed as a percentage, represents the brightness of the color. A lightness of 0% results in black, 50% gives a neither-dark-or-light color, and 100% results in white. The HSL color space facilitates straightforward inversion, as the lightness can be inverted independently from hue and saturation.

The YIQ color model [6] is composed of three channels: Luminance (Y), In-phase (I) and Quadrature (Q). The Y channel represents the brightness of the color, with higher values indicating more brightness. Component I encodes the information regarding the orange-blue color spectrum, and Q encodes the information regarding the purple-green color spectrum. In this colorspace the inversion of the Y channel can be used to invert the ‘lightness’ of the image.

The International Commission on Illumination (CIE) introduced a lightness scale complemented by two chromatic scales to reflect the perceptual behavior of the human visual system. The resulting CIElab colorspace [6], embodies a three-axis structure denoted by L* (Lightness), a* (redness–greenness), and b^*^ (yellowness–blueness). This configuration distinctly segregates chromatic attributes from lightness, rendering the CIElab color space advantageous for inverting the background.

The proper transformation and inversion of color images can be done in image analysis software, including ImageJ [7] and FIJI [8]. However, it requires the installation of software and offers limited flexibility. To improve user friendliness, we generated a web based app that is capable of reversing the background of color images without altering the original colors and without requiring the raw data of the image. We examined the characteristics of various alternative color spaces that can be used to invert the background without altering the hues.

Our application is founded on two principal methodologies. The first employs transformations into different color spaces, isolating color from brightness to maintain the original hues while inverting the background. The second methodology focuses on detecting grayscale values in the RGB image and subsequently inverting or modifying these values.

We integrated these methodologies into an accessible and intuitive web application. This application, compatible with various common image formats, supports four different color spaces, three distinct kernels, and gamma adjustment, requiring only the image itself. It is designed to enhance the presentation of images across a broad spectrum of types (fluorescence images, photos, graphs), thereby improving perception and accessibility to detailed and structural information. Finally, the web app can be used to easily switch images between dark and light themes for presentations, display on the web or publications.

## Methods

### Implementation

The ezReverse web application is developed using Python and employs the Python version of Shiny, for creating interactive web applications. This application also integrates several other Python packages for image processing: numpy for data manipulation, PIL for image reading, skimage for image processing, and colorsys for color space transform. The application’s source code is available on GitHub (https://github.com/Morwey/ezreverse), which also serves as the hub for updates, code revisions, and version release information. This manuscript is based on version release v1.0.0 of the app (https://github.com/Morwey/ezreverse/releases/tag/V1.0.0).

The web app can be accessed at: https://amsterdamstudygroup.shinyapps.io/ezreverse/

Upon launching the ezReverse app (figure 1), users are presented with a pre-loaded example image of cells to demonstrate the effect of the method. The example data is also available on the Github repository (https://github.com/Morwey/ezreverse/tree/main/demo_input).

**Figure 1:**
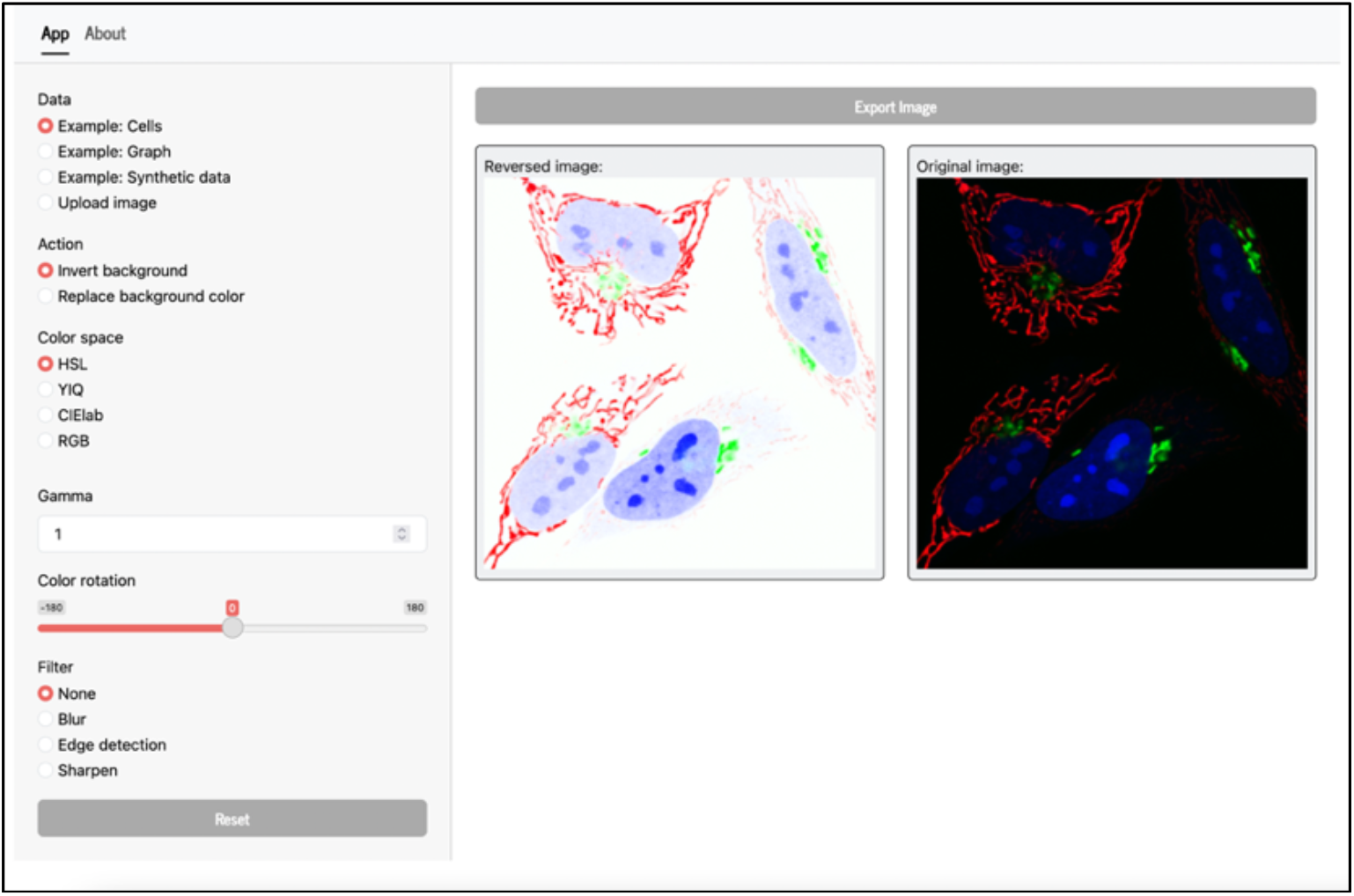
Screenshot of the web app. A panel for user input is located on the left. Users can select example data, the method for the background modification and some other parameters to further tweak the image. On the right, the original image is displayed next to the reversed image and an export button is available for downloading the reversed image.

Three different example images can be selected in the app, which represent a (i) fluorescence image, (ii) a plot of scientific data, and (iii) a synthetic image with a grayscale gradient as a background. These example images can be used to judge the effect of applying the different methods and settings. These example data are used throughout themanuscript to show the results of the different methods and users can therefore readily reproduce the results presented in this manuscript.

A schematic of the processing pipeline is depicted in figure 2 and each of the steps is described in more detail below.

**Figure 2:**
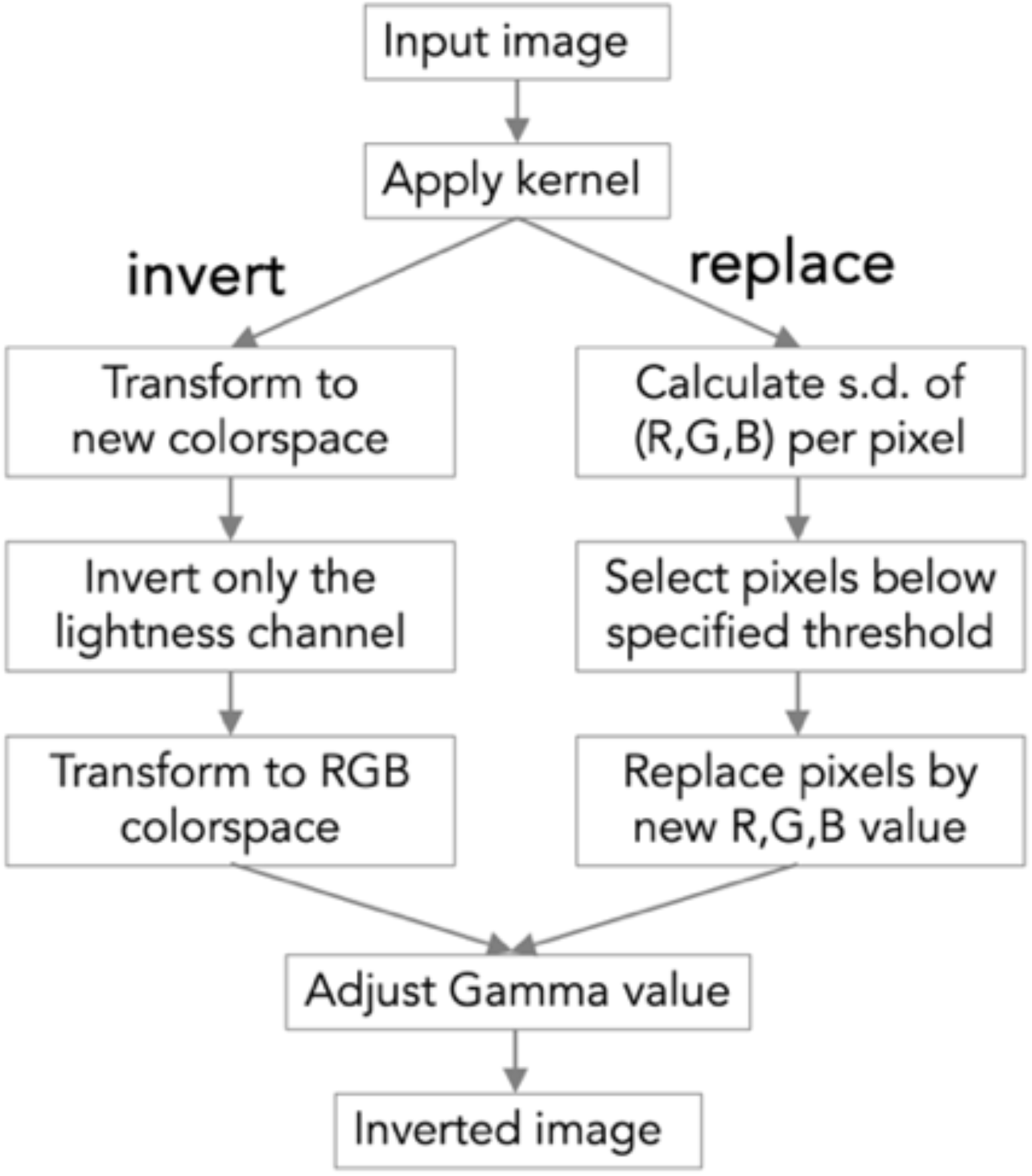
Flow chart that shows the order of the operations for the two methods “invert” and “replace” that are available to reverse the background of the color image.

The ezReverse web application supports a versatile range of image formats for data upload. It can handle any image type that is compatible with the PIL.Image.open method from the Python Imaging Library (PIL), also known as Pillow, including JPEG, PNG, BMP, TIFF, and GIF, among others. After uploading the image, a filter can be applied to smooth or sharpen the image. Next, one of two methods are applied: “invert” background and “replace” background. The invert function operates based on the transformation of various color spaces, while the Background Color Change function is predicated on filtering according to a standard value threshold. These two methods are detailed below.

### Method 1: Invert

Straight inversion of an RGB image changes the background from dark to light, but it also changes the colors, as can be inferred from the images and the pixel distribution in RGB colorspace as shown in figure 3.

**Figure 3:**
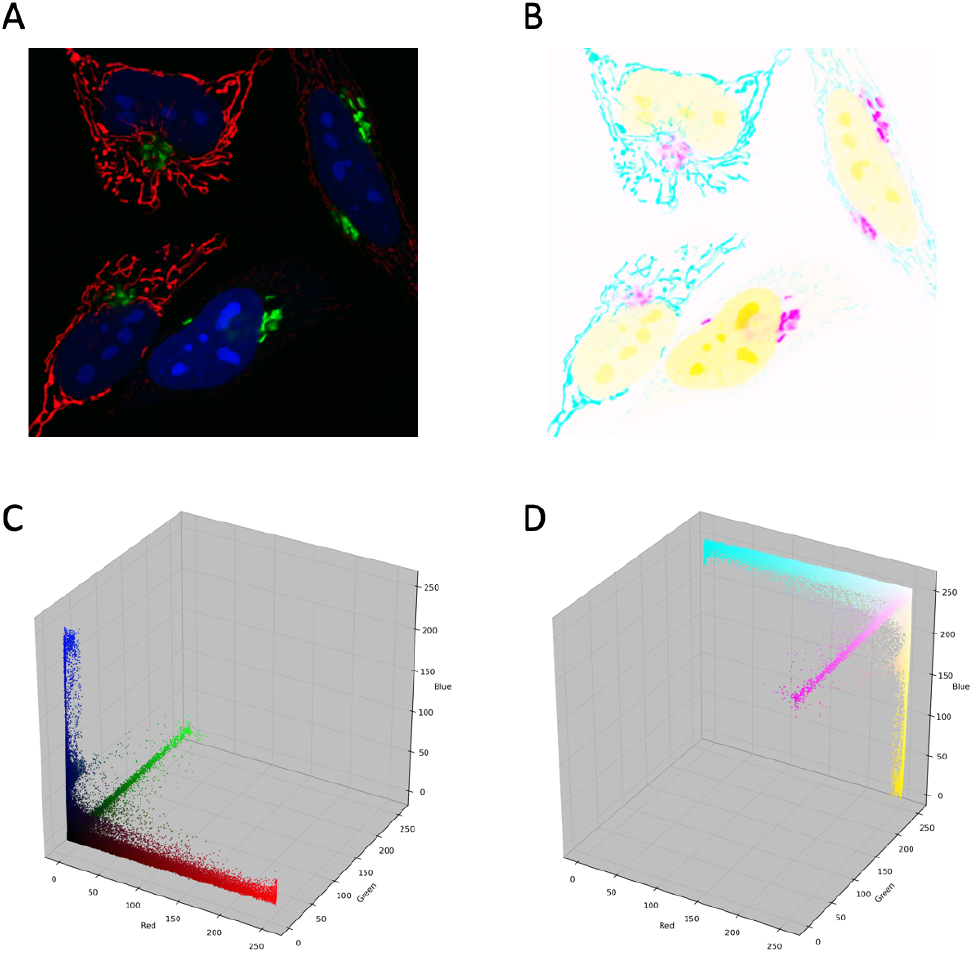
Direct inversion of an RGB image changes the colors. (A) The image from the example data “Cells” that shows fluorescent cells on a black background with red, green and blue hues and (B) the inverted image with a white background. (C) pixel distributions from (A) showing the red, green and blue hues and (D) pixel distribution of the inverted image (B) showing the hues cyan, magenta and yellow.

To retain the original hues, the RGB image can be transformed into other color spaces: CIELab, HLS, and YIQ. Each of these spaces have a channel for ‘lightness’. The app inverts this channel and the resulting image is transformed back to RGB. In figure 4 the transformations are shown in a 3D plot that reflects the pixel distribution in its respective colorspaces. A similar procedure is used for the inversion through the YIQ and CIElab colorspace.

**Figure 4:**
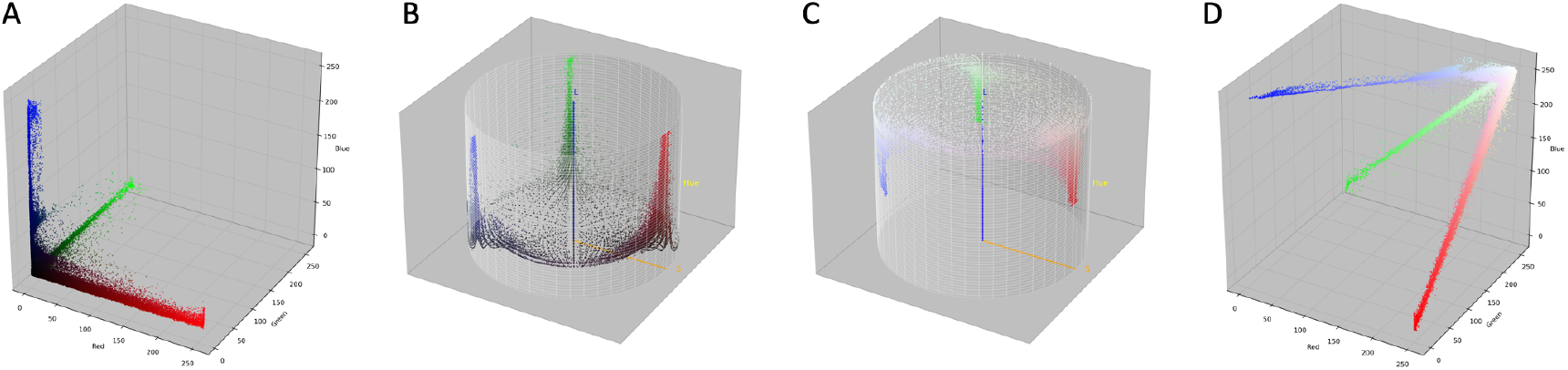
The effect on the pixel distribution in colorspace of the operations that are used to reverse the image. The pixel distribution is based on the example data “Cells” that shows fluorescent cells on a black background with red, green, and blue hues. From left to right: (A) the distribution of pixels from the original image in RGB space, (B) the distribution of the same as (A) transformed into HSL space, (C) the distribution of pixels in HSL space after inverting the L channel (inversion along the z-axis) and (D) distribution of pixels in RGB space after transformation from HSL space, reflecting the distribution in the inverted image.

### Method 2: Replace

This method consists of two steps: identifying gray values and assigning a new color to the identified pixels. The identification method is briefly outlined here. In RGB images, gray values are characterized by similar values for R, G, and B, and so the RGB code for middle gray is 127,127,127. To identify gray values, the standard deviation of the R, G, and B value for each pixel were determined. Pixels with a low standard deviation correspond to gray. In the app, users can set a “tolerance” that defines the maximal standard deviation that is allowed. Subsequently, the pixels that have a standard deviation that is lower than the tolerance are modified. The modification can be inversion (to invert the background), but the gray value pixels can also be replaced by other colors, to allow for more flexibility in the modification of the image.

To further tune the contrast of the image a gamma adjustment is available.

Gamma adjustment utilizes nonlinear methods on the pixels of an input image, thereby altering the saturation of the image. It facilitates a more natural representation of color and luminance, compensating for the nonlinear way human eyes perceive light and color. In the app it can be used to fine-tune the image by modifying the saturation.

### Output

The output image is shown next to the input image to allow direct comparison. Users can directly copy the output image from the browser window or click on the ‘export’ button that will download the result in PNG format.

### Data Visualization

The figures in this manuscript were made with the example data that is included in the ezReverse web app and can be reproduced using the web app. The 3D color distributions were generated in FIJI with the plugin Color Inspector 3D (https://home2.htw-berlin.de/~barthel/ImageJ/ColorInspector/help.htm).

## Use cases

### Method 1: Invert

This method was designed for the inversion of color images, such as those obtained by mutlicolor fluorescence imaging. Here, we show the effect on both a synthetic image with a shaded background, as well as on a 3-color fluorescence image. Figure 5 shows the original input data and the result of the inversion using three colorspaces. Finally, for illustration, the direct inversion in RGB colorspace is also shown. All methods successfully invert the background in both synthetic and fluorescence image. It is clear that the inversion in the colorspaces HSL, YIQ, CIElab, is less destructive than inversion in the RGB colorspace. The HSL colorspace maintains a color closest to the original.

**Figure 5:**
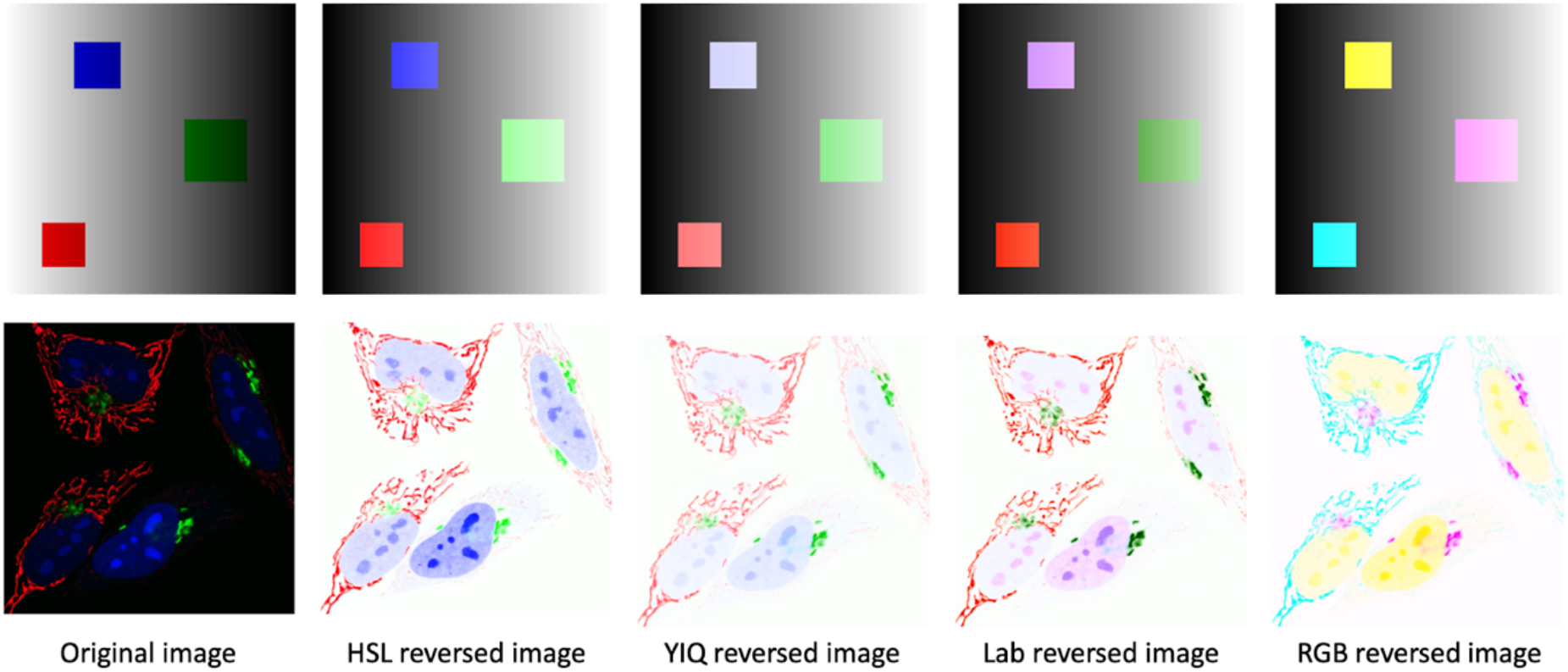
Comparison of Original and inverted images across different color spaces. The upper row showcases synthetic images, while the lower row displays real fluorescent images. From left to right: original image, inverted image employing HSL, YIQ, Lab, and RGB color spaces.

### Method 2: Replace

To validate the effectiveness of the background color change algorithm, we evaluate it through two approaches: utilizing synthetic images and real fluorescence images, with both scenarios set at a threshold of 6.5, and backgrounds replaced by white color. The outcomes are illustrated in figure 6. Here, we also demonstrate the effect of changing the gamma.

**Figure 6:**
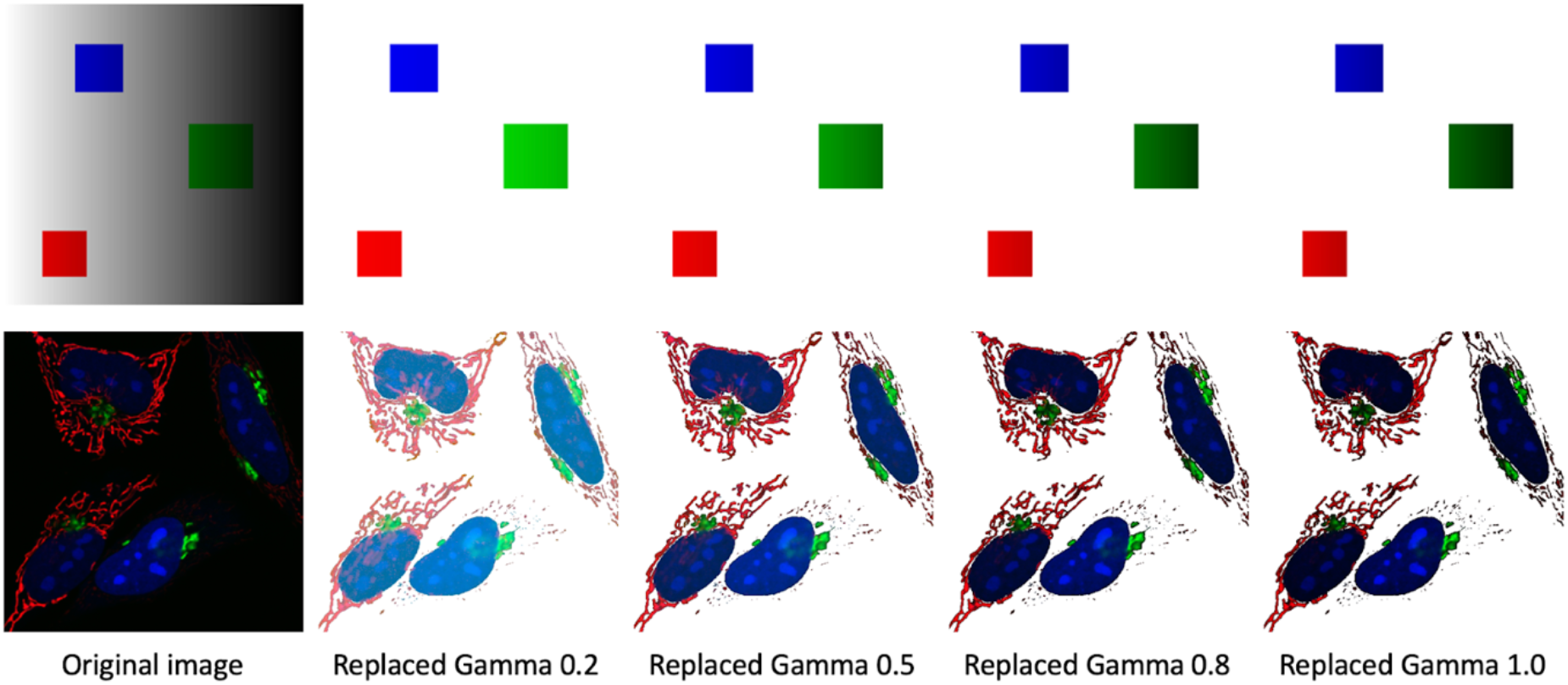
Comparison of Original and Background Color-Altered Images Across Varying gamma Values. The upper row features synthetic images, while the lower row presents real fluorescent images. From left to right: original image, altered image with gamma 0.2, gamma 0.5, gamma 0.8, and gamma 1.0 (default).

The results presented in figure 6 illustrate that the method is effective at replacing the background, both for the synthetic and real fluorescent images. This method uses a threshold to select the gray values, and it works best when there is a clear difference between color and gray values. This is the case for synthetic data. It is less suited for the image with cells, as the difference between color and gray is less clear. Finally, setting the gamma further aids in enhancing the image.

## Discussion

Reversing the background in color images has benefits, but it is not straightforward to do this without changing the colors. To address this issue, we have generated a web based application, ezReverse, that simplifies the manipulation of the background.

We show that the “Invert background” feature works well on both synthetic and real images. Different color spaces yield varied results, the choice of which depends on the use case. Adjusting the suitable gamma value and kernels can enhance the visualization of structures. While the background color change function demonstrates commendable performance on synthetic images, for real images, fine-tuning with the gamma value should yield improved results.

The “replace background” method works well when there is a clear distinction between graylevels (including black and white) and color. The advantage of this method is that it maintains hues identical to the original, facilitating improved visual clarity. However, when the boundaries between color and greys is subtle (as for fluorescence images), the “invert” method may be preferrable.

Compared to existing methods offered by image processing software, e.g. ImageJ, the ezReverse web app exhibits greater ease of use and provides parameters for fine-tuning. Its deployment on an online platform facilitates its use and access without the necessity for installation. Additionally, it accommodates a wide variety of image types. Altogether, the ezReverse web app is user-friendly and allows for basic customization of the visualization.

### Limitations

The advantage of a web app is its accessibility and the platform and software independent operation, but this comes with the limitation that an internet connection is required. Since images can be large, the speed partially depends on the bandwidth of the connection. Another limitation is that the current app does not accept movies. Finally, the inversion of the lightness channel uses a simple, linear method and other, more sophisticated methods may improve the quality of the inverted image.

### Outlook

The shiny package was originally developed to generate web apps that are based on R-code. Recently shiny was extended to Python. Since Python has many libraries for (bio)image analysis and deep learning, the option to generate web apps with Shiny for Python increases potential applications. We hope that our app is an example of how Python can be combined with Shiny and that this will be an inspiration for new web apps.

## Acknowledgments

The tweets of Christophe Leterrier promoting and explaining the use of the #invertedLUT (e.g. https://twitter.com/christlet/status/921763282893070336) were a significant (p<0.05) source of inspiration. The discussions on the image.sc forum has motivated us to come up with a web app that uses some of the proposed methods (https://forum.image.sc/t/invert-rgb-image-without-changing-colors/33571).

## Data availability

All data and code is publicly available: https://github.com/Morwey/ezreverse

## Author contributions

X.S. designed the app, wrote the code and wrote the manuscript; J.G. designed the app, and wrote the manuscript.

## References

1. Specht EA, Braselmann E, Palmer AE. A Critical and Comparative Review of Fluorescent Tools for Live-Cell Imaging. Annual Review of Physiology. 2017;79: 93–117. doi:10.1146/annurev-physiol-022516-034055

2. Sanderson MJ, Smith I, Parker I, Bootman MD. Fluorescence Microscopy. Cold Spring Harb Protoc. 2014;2014: pdb.top071795. doi:10.1101/pdb.top071795

3. Johnson J. Not seeing is not believing: improving the visibility of your fluorescence images. MBoC. 2012;23: 754–757. doi:10.1091/mbc.e11-09-0824

4. Altunkeser A, Körez MK. Usefulness of grayscale inverted images in addition to standard images in digital mammography. BMC Medical Imaging. 2017;17: 26. doi:10.1186/s12880-017-0196-6

5. Park JB, Cho YS, Choi HJ. Diagnostic accuracy of the inverted grayscale rib series for detection of rib fracture in minor chest trauma. The American Journal of Emergency Medicine. 2015;33: 548–552. doi:10.1016/j.ajem.2015.01.015

6. Ramanath R, Drew MS. Color Spaces. In: Ikeuchi K, editor. Computer Vision: A Reference Guide. Cham: Springer International Publishing; 2021. pp. 184–194. doi:10.1007/978-3-030-63416-2_452

7. Schneider CA, Rasband WS, Eliceiri KW. NIH Image to ImageJ: 25 years of image analysis. Nature Methods. 2012;9: 671–675. doi:10.1038/nmeth.2089

8. Schindelin J, Arganda-Carreras I, Frise E, Kaynig V, Longair M, Pietzsch T, et al. Fiji: an open-source platform for biological-image analysis. Nature Methods. 2012;9: 676–682. doi:10.1038/nmeth.2019

